# Inferring adaptive gene-flow in recent African history

**DOI:** 10.1101/205252

**Authors:** George Busby, Ryan Christ, Gavin Band, Ellen Leffler, Quang Si Le, Kirk Rockett, Dominic Kwiatkowski, Chris Spencer

## Abstract

Gene-flow from an ancestrally differentiated group has been shown to be a powerful source of selectively advantageous variants. To understand whether recent gene-flow may have contributed to adaptation among humans in sub-Saharan Africa, we applied a novel method to identify deviations in ancestry inferred from genome-wide data in 48 populations. Among the signals of ancestry deviation that we find in the Fula, an historically pastoralist ethnic group from the Gambia, are the region that encodes the lactose persistence phenotype, *LCT/MCM6*, which has the highest proportion of Eurasian ancestry in the genome. The region with the lowest proportion of non-African ancestry is across *DARC*, which encodes the Duffy null phenotype and is protective for *Plasmodium vivax* malaria. In the Jola from the Gambia and a Khoesan speaking group from Namibia we find multiple regions with inferred ancestry deviation including the Major Histocompatibility Complex. Our analysis shows the potential for adaptive gene-flow in recent human history.

## Introduction

The source of adaptive genetic variation remains an important question in evolutionary biology (Hedrick, 2013). In the canonical view of evolution, natural selection causes a change in the frequency of a mutation that was either present before the introduction of selection, so-called standing variation, or a new mutation that arose after the selective pressure began (Hermisson and Pennings, 2005). A third potential source of novel genetic variation on which natural selection can act is via gene-flow resulting from an inter or intra species admixture event. Examples of such adaptive introgression include the exchange of mimicry adaptations amongst *Heliconius* butterfly species (Consortium, 2012), the transfer of insecticide resistance genes between sibling *Anopheles* mosquito species (Clarkson et al., 2014), and the spread of pesticide resistance across populations of house mice (Staubach et al., 2012).

In the human lineage, recent genomic studies have shown that our demographic history is complex, involving population merges as well as splits (Green et al., 2010; Patterson et al., 2012; Prüfer et al., 2014; Hellenthal et al., 2014; Fu et al., 2015). These merges, which are usually referred to as admixture or introgression events, open the door to the transfer of advantageous alleles. For example, we now know that trysts between archaic *Homo sapiens* and Neanderthals over fifty thousand years ago (kya) transferred beneficial alleles from Neanderthals into the human lineage (Green et al., 2010; Patterson et al., 2012; Prüfer et al., 2014; Fu et al., 2015; Abi-Rached et al., 2011; Gittelman et al., 2016). While genomic regions with elevated archaic ancestry have been observed, it is also possible to identify regions that are devoid of archaic ancestry and where it is thought that archaic alleles at these loci have been at a disadvantage (Sankararaman et al., 2014; Racimo et al., 2015).

Despite the prevalence of more recent admixture events over the past 50 kya of human history, relatively few instances of adaptation from these events have been found. Notable known examples include the exchange of high altitude adaptations between the ancestors of Sherpa and Tibetans (Jeong et al., 2014) and the spread of the *Plasmodium vivax* malaria-protecting Duffy null mutation in Madagascar as result of gene-flow from mainland Bantu-speaking Africans (Hodgson et al., 2014).

Within Africa, multiple studies using different techniques have shown admixture to be a common theme in the recent history of the continent Pagani et al. (2012); Schlebusch et al. (2012); Pickrell et al. (2012, 2014); Pickrell and Reich (2014); Busby et al. (2016); Patin et al. (2017); Carlton and Sullivan (2017); Amer (2017); Skoglund et al. (2017). As a result, the genomes of individuals from these populations contain segments which derive from multiple ancestries. Inferring whether gene-flow has assisted adaptation requires inferring how these ancestries change locally along the genome. Although a number of strategies exist for local ancestry inference (Price et al., 2009; Baran et al., 2012; Brisbin et al., 2012; Maples et al., 2013), most are designed to distinguish at best continental-level ancestry from a small number of reference populations.

The approach we develop here for inferring whether gene-flow has contributed more or less ancestry than expected at a locus involves sampling from a Hidden Markov Model to identify the likely donor haplotype from a large set of reference genomes. We use this chromosome painting approach to assign a donor ancestry label (which can be at individual, population, region, or continental level) at each locus in the genome to all recipient individuals. Iterating this process incorporates ancestry assignment uncertainty and we infer the ancestry proportions from different donor groups across the genome.

We use this analysis to ask a simple question: across individuals in these admixed populations, are there regions of the genome where ancestry is significantly deviated away from genome-wide expectations? We construct a statistical model to test for significant deviations and interpret deviations in local ancestry as possibly resulting from natural selection, which can act either by increasing the frequency of the introgressed haplotype (positive selection) or prevent it from replacing an established haplotype (negative selection). We highlight specific examples where ancestry deviations align with known targets of selection, describe patterns of selection across the data, and discuss the challenges in using approaches of this kind for detecting adaptive gene-flow.

## Results and Discussion

### Inferring local ancestry

We used a published dataset containing computationally phased haplotypes from 3,283 individuals, from 60 worldwide populations, typed at 328,176 high quality genome-wide SNPs [Fig. S1] (Busby et al., 2016) and grouped individuals from these populations into 8 separate ancestry regions, based on genetic and ethno-linguistic similarity, as described previously (Busby et al., 2016) [Fig. 1a]. We painted chromosomes from these populations using *ChromoPainter* with donors which did not include any closely related populations, here defined as individuals from within their own ancestry region. Therefore, for each haplotype, there are seven non-local ancestries (or colours of paint) from which they can copy.

**Figure 1:**
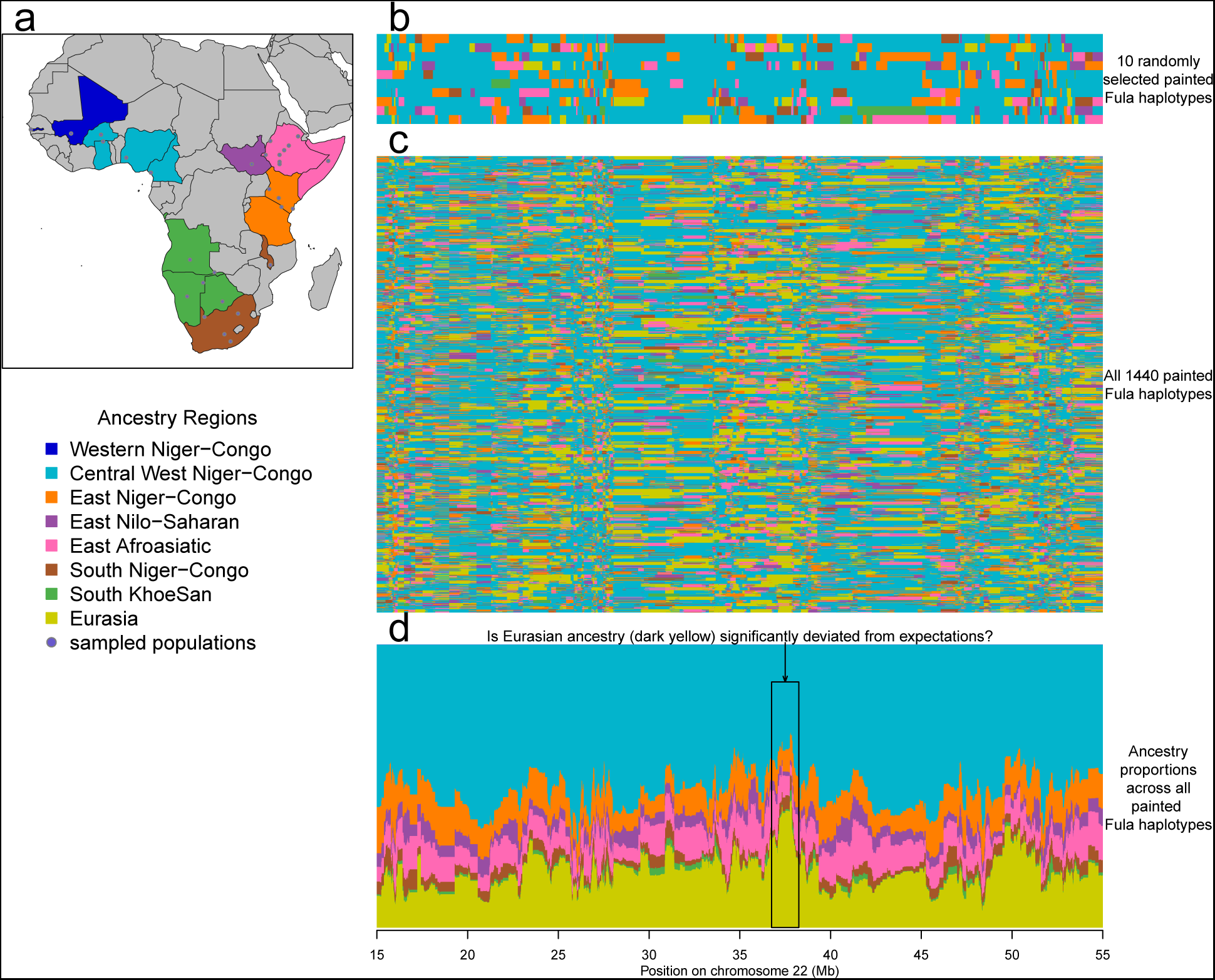
Overview of local an ces try based analysis. **a** Location of the 48 populations from seven Africa n ancestry regions used in the analysis; **b** chromosome painting samples (rows) of chromosome 22 (3392 SNPs) from ten Fula haplotypes show mosaic local ancestry; c chromosome painting samples from all 144 Fula haplotypes, each set of 10 consecutive rows are paintings from one recipient haplotype; **d** chromosome painting samples from (c) stacked to show ancestry proportions locally along chromosome 22.

### Identifying changes in local ancestry

Our approach for inferring ancestry deviations is based on the idea that ancestry tracts shared between two populations will reveal a signal of admixture (Hellenthal et al., 2014), but will be randomly distributed amongst the genomes of individuals within those populations (Baird, 2006)[Fig. 1b]. So, whilst individuals will have mosaic (i.e. block-like) ancestry [Fig. 1c], when summed across all individuals in a population, ancestry proportions at specific loci (SNPs) will resemble genome-wide proportions in expectation [Fig. 1d]. In this framework, we would expect ancestry proportions at a SNP to match genome-wide proportions, with some variation which we can model. A significant deviation away from expected ancestry proportions suggests that natural selection may have contributed to the change. We note that this will only ever allow for indirect inference of selection. This is nevertheless similar in spirit to other commonly used selection identification methods, such as the integrated haplotype score (iHS) (Sabeti et al., 2002; Voight et al., 2006) which scans the genomes of a population for longer than expected haplotypes based on their frequency, which are then (indirectly) inferred to have swept as a result of selection.

To help account for the uncertainty in ancestry inference we sampled chromosome paintings 10 times for each individual. We use a binomial likelihood to test whether the ancestry at each SNP along the genome differs from expectations. Specifically, for each recipient haplotype, we infer the expected proportion of ancestry from each non-self ancestry region from their genome-wide ancestry paintings. At each position, we fit a deviation parameter, *β*, that best explains the sampled ancestries at that SNP across all individuals. *β* can be thought of as the increase (or decrease) in expected ancestry proportions across individuals on the logit scale that best explains the proportions observed at a SNP compared to the genome-wide average. To generate a *P* value, we compared the likelihood ratio test statistic to a *χ*^2^ distribution. At each SNP in the genome, this test gives a *P* value and estimate of *β* for each of the 7 non-self ancestries tested.

Modelling ancestry in this way has the advantage that it adjusts for an individual’s specific ancestry proportions. The likelihood is a product across individuals which assumes that individuals are independent given the underlying ancestry proportions. Quantile-quantile (Q-Q plots) of the observed test statistics typically show high levels of inflation [Fig.]S2. There are at least three explanations for this inflation. Firstly, due to shared ancestral histories among haplotypes from the same population, the paintings are unlikely to be independent between samples in a population, so the assumption in the likelihood that the samples are independent is likely to be false. Moreover the demographic process that has given rise to the ancestry paintings is more complex than the model, leading to local variations in paintings that are not captured by the null. Secondly, the chromosome painting model explicitly uses correlations in ancestry locally in the genome (i.e. linkage disequilibrium) meaning that tests are not independent along the genome, as shown by the autocorrelation plots in Figure S2. Finally, adaptive introgression may have a weak effect genome-wide. We discuss below alternative approaches that model the variance (as well as the expectation) in ancestry proportions across the genome. However, these limitations mean that it is difficult to estimate a genome-wide threshold for significance in the traditional sense (i.e. for defining false positive/negative rates), so we interpret *P* values as providing a relative ranking rather than an absolute measure of significance.

### Ancestry deviations in the Fula identify known targets of selection

We illustrate our approach with an analysis of genomes from the Fula ethnic group from The Gambia. The Fula (also known as Fulani) are historically nomadic pastoralists spread across West Africa who we inferred in our previous analysis to have experienced an admixture event involving a largely Eurasian source (0.19 admixture proportion) mixing with a West African source around 1,800 years ago (239CE (95% CI = 199BCE-486CE)). (Admixture events across all analysis populations inferred previously (Busby et al., 2016) are summarised in Figure S3). This event introduced a significant amount of Eurasian ancestry into the Fula which is relatively easy to identify given that Eurasian populations split and drifted from African populations over 50kya (Mallick et al., 2016). They thus have haplotypes that are easier to differentiate from each other, making our approach relatively well powered to detect deviations in ancestry proportions.

In the Fula, the region of the genome with the lowest proportion of Eurasian ancestry (−*log*_10_*P* = 10, *β*=-4 across all Fula individuals) is on chromosome 1 and contains the Duffy Antigen Receptor *DARC* gene [Fig. 2a] whilst the region with the highest level of Eurasian ancestry (−*log*_10_*P* = 15, *β*=2) is on chromosome 2 and contains the *LCT* and *MCM6* genes [Fig. 2b]. Polymorphisms in *DARC* form the basis of the Duffy blood group system, and the Duffy null mutation, which provides resistance to *Plasmodium vivax* malaria (Miller et al., 1976) is almost fixed across Africa and absent outside (Howes et al., 2011). Our observation less Eurasian than expected at this locus suggests that African haplotypes have been beneficial at this locus after the admixture event that brought the Eurasian ancestry into the Fula; that is, within the last 2,000 years. Similar effects at this locus have been observed before: based on an analysis of allele frequencies, selection following admixture between Austronesians and sub-Saharan Bantus is likely to have driven frequencies of the Duffy null mutation to higher than expectations based on admixture proportions in the Malagasy of Madagascar (Hodgson et al., 2014), and a recent study of Sahel populations showed an excess of African ancestry at this locus in Sudanese Arabs and Nubians (Triska et al., 2015).

**Figure 2:**
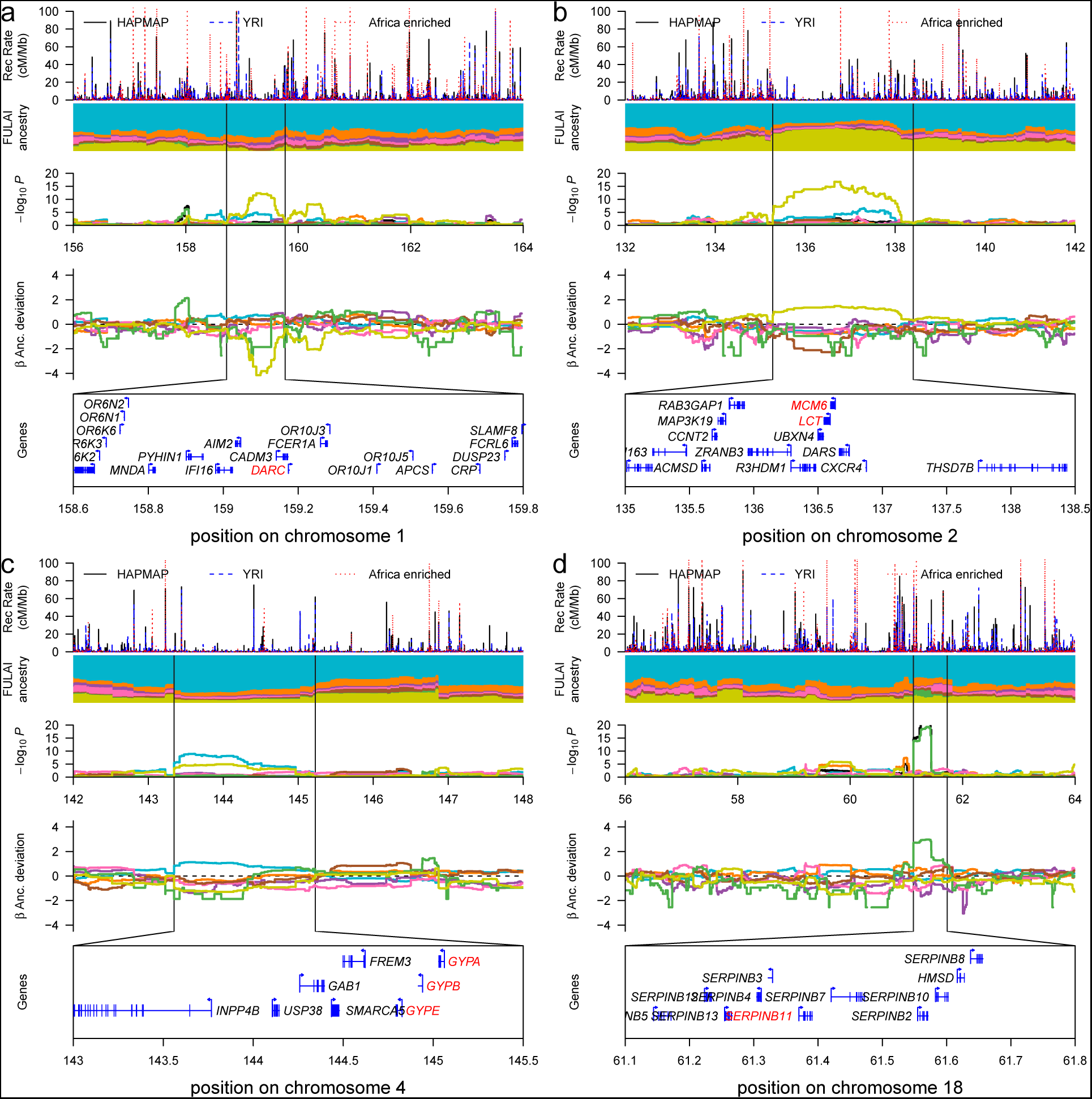
Local ancestry deviations in the West African Fula. Each main panel is made up of 5 sub-panels showing, from top to bottom: the recombination rate across the region; ancestry proportions coloured by region as in Figure 1; evidence for deviation based on the binomial deviatio n model; *P* values for the likelihood ratio test, coloured by ance stry; the associated *β* coe fficient fit by the LRT for each ancestry; the genes in the region. **a** lack of Eurasian ancestry around *DARC* on chromoso me 1; **b** excess Eurasian ancestry around the *LCT* and *MCM6* genes on chromosome 2; c increased Central West African ancestry at the GYP genes on chromosome 4. **d** increased Khoesan copying at the SERPIN genes on chromosome 18

Mutations in an intron in *MCM6*, a regulatory element for *LCT* lead to the lactose persistence phenotype (Ingram et al., 2008) (the ability to digest milk into adulthood), and represent one of the clearest signals of recent natural selection in the genome. Lactose persistence has evolved at least twice independently (Bersaglieri et al., 2004), in Europe and in East Africa (Ranciaro et al., 2014). Encouragingly, our analysis identifies this Eurasian haplotype, despite the SNP set not containing the ‘European’ lactase variant [13910 C>T polymorphism, rs4988235; Fig. S4] and suggests that the European haplotype has entered the Fula as a result of gene-flow from Europe approximately 2kya. In both this case and that of *DARC* described above, our ancestry based analysis provides new insight into the potential origins of selected mutations.

### The genome-wide distribution of ancestry deviations in the Fula

We note that because Eurasian and Khoesan haplotypes are likely to be more diverged from those of other sub-Saharan African groups we are better able to characterise them and so have more power to identify deviations concerning these ancestries. Indeed, over half (21) of the top 40 ancestry deviations observed in the Fula involve Eurasian ancestry [Table S1; Fig. S5], including increases of Eurasian ancestry across the *HLA* region on chromosome 6 (−*log*_10_*P* = 12.38, *β* = 1.29), a group of *DSC* genes on chromosome 18 (−*log*_10_*P* = 13.55, *β* = 1.35), and at *APOL* genes on chromosome 22 (−*log*_10_*P* = 13.01, *β* = 1.32). Of the non-Eurasian signals, of note are a significant increase in Central West African ancestry across the *GYP* genes on chromosome 4 [Fig. 2c] (−*log*_10_*P* = 8.96, *β* = 1.13), encompassing a region which has recently been identified as being associated with malaria (Malaria Genomic Epidemiology Network, 2015; Leffler et al., 2016), and which, based on a recent iHS analysis (Johnson and Voight, 2017), shows signs of being under natural selection. We observe an increase in Khoesan ancestry across the *SERPIN* genes on chromosome 18 [Fig. 2d] (−*log*_10_*P* = 19.32, *β* = 2.98), one of which, *SERPINB11*, has previously been implicated as a target of natural selection in pastoralist groups in Africa (Seixas et al., 2012).

We next looked at the evidence for covariance in ancestry deviation at SNPs across the genome. As expected, in all cases we found negative covariance in ancestry at SNPs where we observed a significant deviation signal [Fig. 3a]. When one ancestry increases in frequency at a locus, it must be at the detriment to other ancestries. This association was more pronounced for more closely related ancestries. For example, the largest negative relationships were seen between East African groups [red boxes in Fig. 3a]. Interestingly, when we looked at the distribution of *β*s at significant SNPs, there was a general trend towards signals of ancestry increases, except in the case of Eurasian (yellow) and Central West African (light blue) ancestry [Fig. 3c].

**Figure 3:**
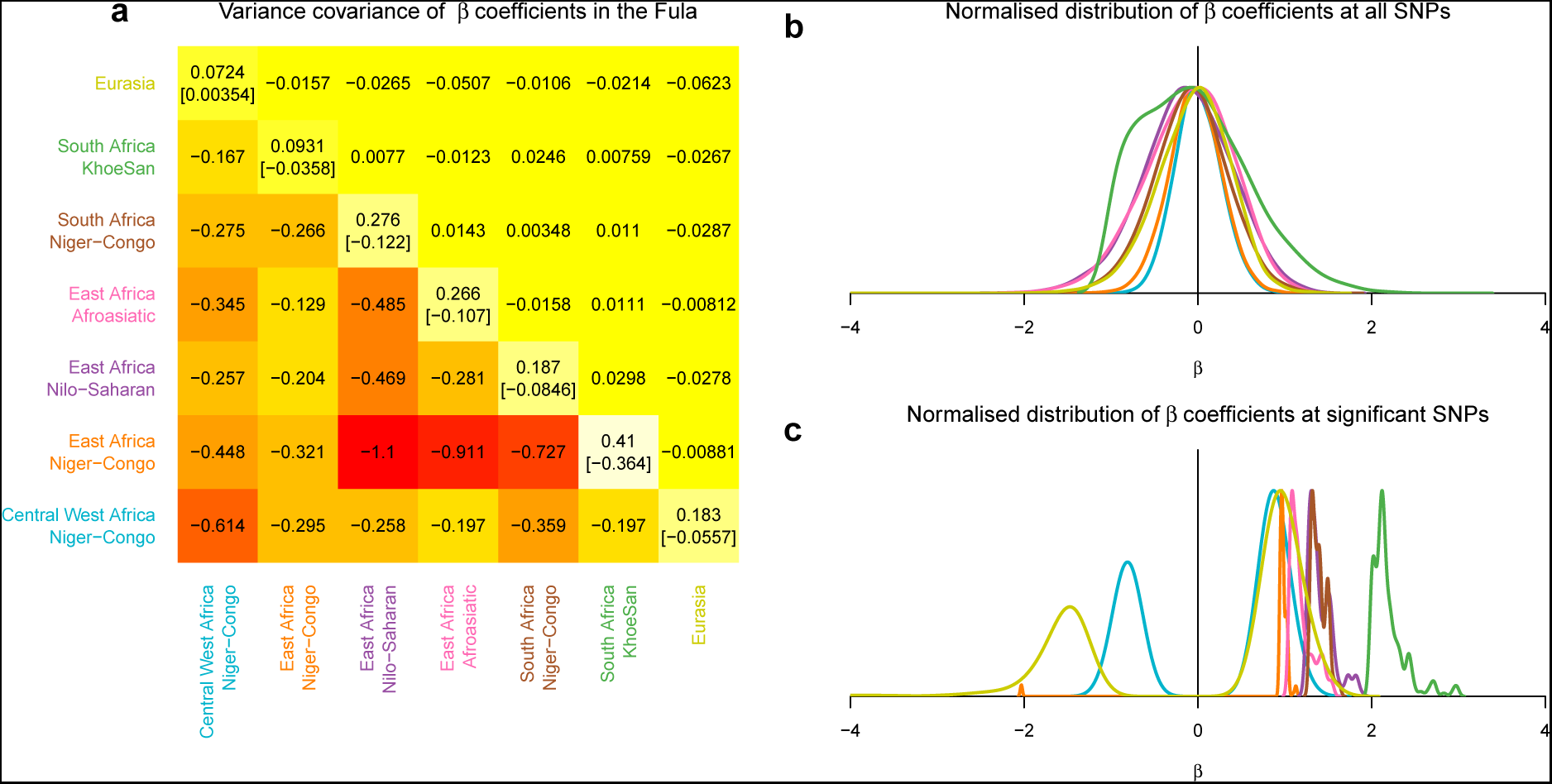
Negative covariance of ancestry deviations in the Fula. **a** top right diagonal shows of heatmap shows covariance at genome-wide SNPs; bottom left are SNPs that are significant in either of the ancestry region pair; the diagonal shows the variance and mean in brackets within each ancestry region **b** the genome-wide distribution of *β* coefficients, split by ancestry, across all S NPs and normalised to have a maximum density of 1. **c** the distribution of *β* coefficients for significant SNPs, normalised to have a maximum density of 1. in panels **b** and c lines are coloured by ancestry as in panel a

### Accounting for reciprocal local ancestry deviations

One concern with our analysis is that deviations in local ancestry might result from reciprocal shared ancestry. That is, a non-local donor individual might share haplotypes with a recipient population because they themselves are admixed from the recipient group. We will observe such reciprocal copying in our framework when a single, or small number of, donor individuals contribute excessively to an ancestry deviation signal. An example can be seen in Figure 4. Across all Gambian populations from West Africa (of which we show the Wollof as an example) we observed a significant increase in East African ancestry across the 32.2-33Mb region of chromosome 6, which contains several *HLA* genes [Fig. 4a and 4b]. However, this signal is driven almost entirely by Wollof individuals copying from a single Nilo-Saharan speaking individual, Anuak11. When we painted this individual with both non-local and local donors (i.e. including individuals from the Nilo-Saharan ancestry region), across this region we observe that one haplotype from this individual copies a roughly 1Mb chunk of their genome from the West African ancestry region that includes Gambian populations [dark blue in Fig. 4c] even though there were donors from the more closely related Nilo-Saharan ancestry region available. Our analysis of admixture in the Anuak showed an admixture event involving Central West African and Nilo-Saharan sources in 703CE (95% CI 427CE-1037CE) [Fig. S3], which may have provided a vehicle for West African ancestry to enter this population.

**Figure 4:**
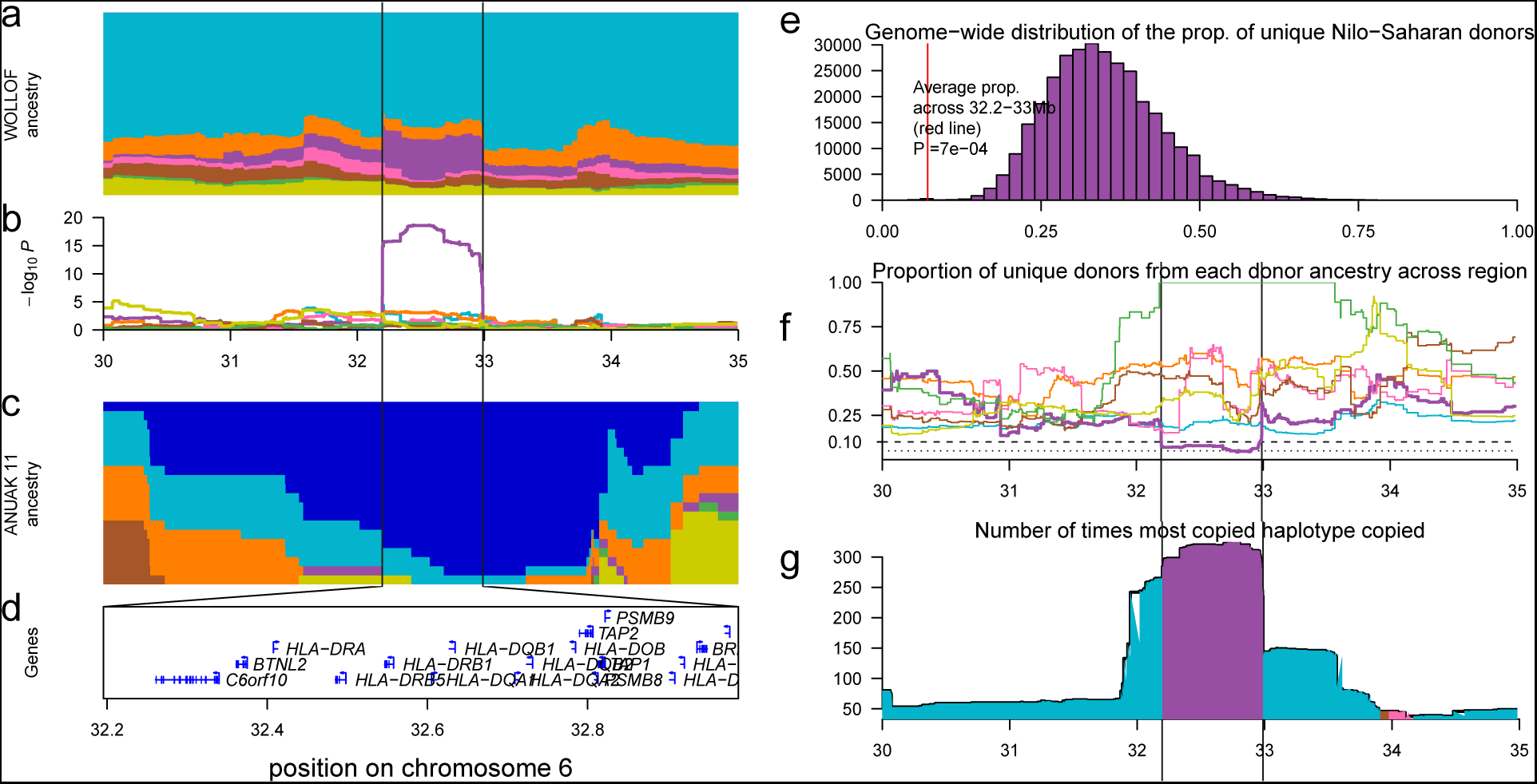
Investigating local ancestry at the ULA in the Wollof. **a** local ancestry across 2160 Wollof haplotypes at the 30-35Mb region of chromosome 6. **b** results of the binomial ancestry deviation test shows a significant increase in Nilo-Saharan (purple) ancestry between 32.2 and 33Mb. c ances try proportions from a local painting analysis in a single Anuak individual showing Western African (dark blue) ancestry. **d** UCSC genes across this region. **e** genome-wide distribution of unique Nilo-Saharan donors in the Wollof. **f** the proportion of unique donors when each of 7 non-self ancestries are copied across the 30-35Mb region of chromosome 6. g the numberof times the most common haplotype is copied; the colour under the line represents the ancestry of the most copied individual.

One way to quantify this effect is to look at the number of unique donor haplotypes from an ancestry region contributing to a signal as a proportion of the total number of donors. This quantity will be 1 if every recipient individual copies from a different donor haplotype and will tend towards zero as the number of unique donors copied from goes down. Whilst for Nilo-Saharan ancestry in the Wollof the median proportion of unique donors contributing to a signal genome-wide is 0.342 (95%CI = 0.20-0.55) [Fig. 4e], the average across the 32.2-33.Mb region of chromosome 6 is 0.072, which lies significantly outside of the genome-wide distribution (empirical outlying *P* value = 0.0007). We can see that this signal is driven by a low proportion of unique donors from the Nilo-Saharan ancestry region [purple line in Fig. 4f] and a large increase in copying from the most copied haplotype [Fig. 4g]. Because this effect may be the result of the donor individual being admixed from the recipient population, in such cases, it is not possible to infer the directionality of ancestry sharing. Therefore, to guard against reciprocal copying we preformed an additional filter on the results. At each locus in the genome, we computed the number of unique donor haplotypes that contributed to the total amount of copying from that region. If the number of unique donors was ≤ 3 or the proportion of unique donors was < 0.1, we assumed the result was due to reciprocal copying.

### Patterns of ancestry deviation across populations

To explore ancestry deviations across the full dataset, we generated a table of the top 159 signals across all populations. These 159 signals were all above a conservative significance threshold of *-log*_10_*P* ≥ 8 [Fig. 5; Table S1], although we note again identifying the correct threshold for our approach is difficult. In just under two thirds (31) of the other 46 sub-Saharan African populations we observed no significant genome-wide deviations in ancestry. We observed multiple (> 5) significant signals in just 3 populations: the Jola and Fula from the Gambia, and the Ju/hoansi from Namibia. Around a quarter (39/159; 24.5%) of the loci identified across the 48 populations, of which 32 were in the Jola, Fula and Ju/hoansi, were discounted from further analysis because of potential reciprocal copying, resulting in a final list of 114 loci, 79% of which were in the Fula, Jola, and Ju/hoansi.

**Figure 5:**
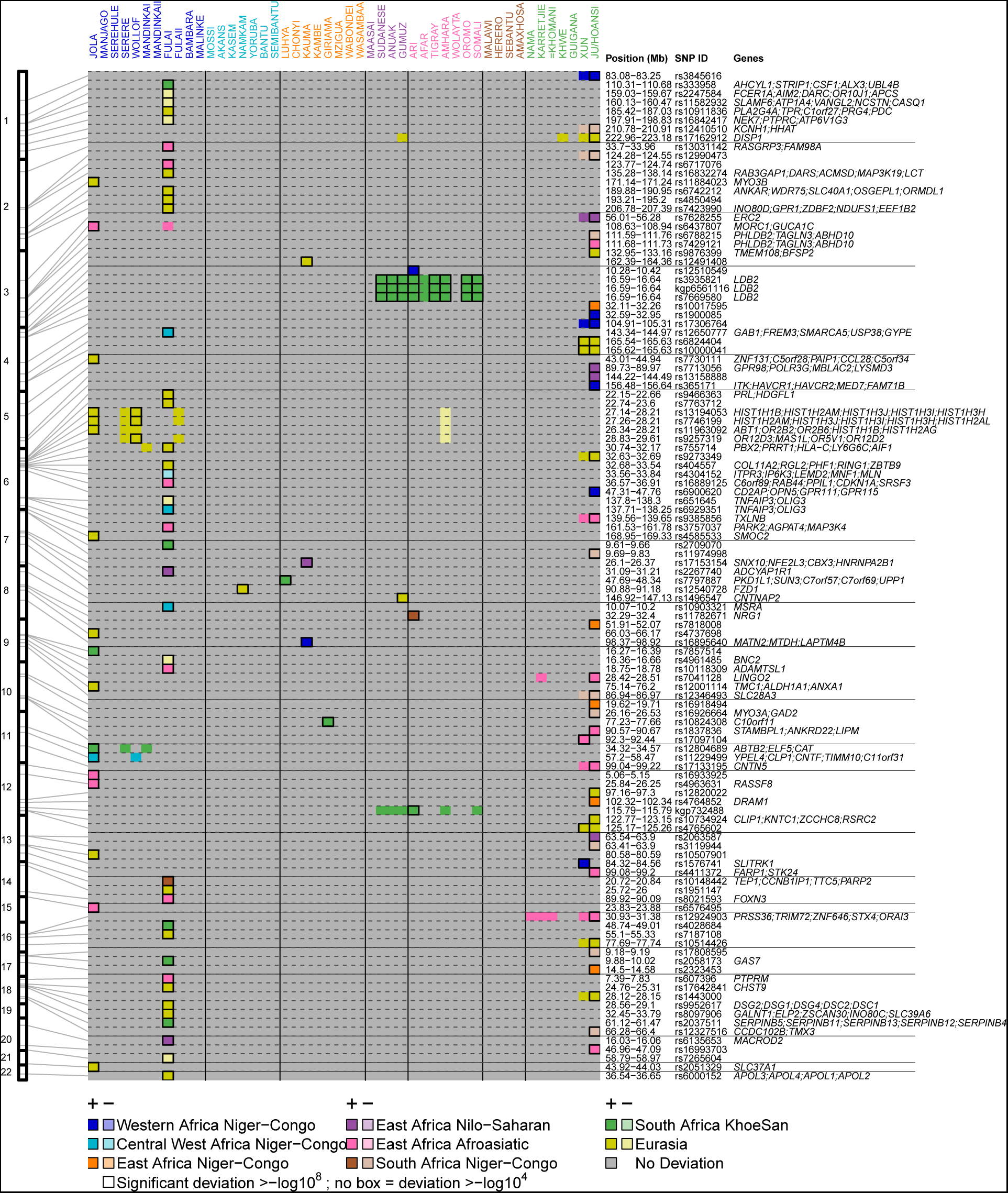
Genome-wide ancestry deviations across sub-Saharan Africa. For each of the 114 loci (rows) where we observe ancestry deviations with a *−log*_10_*P* ≥ 8, for each population (columns) we show the ancestry that has changed (colour) and the direction of change (lighter colours show decrease, darker increase) and the strength of significance (black box 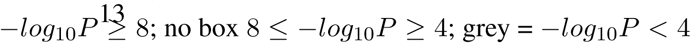. The positions of the significant loci are shown on the right and genes of interest, together with the detailed genomic region, are shown on the left.

As mentioned, in addition to the Fula, we observed multiple ancestry deviations in another Gambian population the Jola [Fig. 5; Fig. S6]. In this group the most deviated region of the genome involves a significant increase in Eurasian ancestry at 27.3-28.2Mb on chromosome 6 which containing several *HLA* genes (*−log*_10_*P* = 12.76, *β* = 1.94), a signal that was also observed in the Wollof, also from the Gambia (*−log*_10_*P* = 10.41, *β* = 1.57). We also observed a significant increases of Eurasian ancestry at *MYO3B* (*−log*_10_*P* = 13.53, *β* = 1.97) and *SMOC2* (*−log*_10_*P* = 9.40, *β* = 1.75), and an increase in Khoesan ancestry on chromosome 11 at *NLRP10* (*−log*_10_*P* = 12.87, *β* = 2.24), which codes for a protein involved in inflammation and apoptosis (Inohara and Nunñez, 2003; Tschopp et al., 2003).

In the Ju/hoansi, the top region of differentiation in the genome involves an increase in East Africa Niger-Congo ancestry at 26.1-26.6Mb on chromosome 10 (*−log*_10_*P* = 17.14, *β* = 2.07) which contains *MYO3A*, a gene involved in hearing (Walsh et al., 2002), and *GAD2*, which has been associated with decreased body mass index in Europeans (Boesgaard et al., 2007). We also observe a significant increase (*−log*_10_*P* = 11.48, *β* = 1.86) of East Africa Afroasiatic ancestry at 99.08-99.20Mb on chromosome 13, a region containing *FARP1* and *STK24*. *FARP1* is thought to be a member of the band 4.1 protein superfamily (Koyano et al., 1997), a major structural component of red blood cell membranes (Conboy et al., 1986). We also see an increase in West African Niger-Congo ancestry at 47.65-47.76Mb on chromosome 6 a region containing *CD2AP*, *OPN5*, *GPR111*, and *GPR115* genes, and a large increase in Eurasian ancestry at *DISP1* on chromosome 1 (*−log*_10_*P* = 10.70, *β* = 2.88).

Eight populations [Fig. 5] showed evidence of significant ancestry deviation at a short haplotype in the middle of the *LDB2* gene on chromosome 4. All 8 of these populations come from East Africa and are either Nilo-Saharan or Afroasiatic speakers [Table S1; Fig. S8]. We sought to characterise the nature of this signal further by combining data across all 235 individuals from these 8 populations (Sudanese, Somali, Anuak, Gumuz, Ari, Tigray, Amhara and Oromo) and the two other populations from the region with nominal evidence of deviation (Afar and Wolayta). In all cases, we observed a signal of increased Khoesan ancestry [Fig. S8a] and localised this to increased copying specifically from the Ju/hoansi, Xun, and Amaxhosa of southern Africa [Fig. S8d]. This differs considerably from a randomly selected SNP 1.5Mb downstream, with a more usual copying profile and where the Luhya and several Eurasian groups were typically copied more. *LDB2* codes for a LIM-domain binding protein, *LDB2*, which is a transcriptional regulator and has been linked to carotid atheroschlerosis (Mm et al., 2014).

### Considerations of our approach to infer adaptive gene-flow

Whilst admixture appears to be an almost universal characteristic of the populations that we use in this analysis, we only find evidence of substantial adaptive gene-flow in three populations: the Fula and Jola from the Gambia and the Ju/hoansi from Namibia. Importantly, we infer significant recent admixture involving diverged source populations in these three groups, from Eurasian-like sources in the Gambians, and East African Asiatic-like source in the Ju/hoansi [Fig. S3] (Busby et al., 2016). Our method is likely to have greater power to detect deviations in these scenarios. As such, it is perhaps not surprising that across populations from the Central West African ancestry region, where we previously inferred admixture events involving mostly closely related African sources (Busby et al., 2016) we infer a single example of adaptive gene-flow in the Namkam from Ghana at *FZD1* on chromosome 8 [Fig. 5]. It is important to note that even though all Gambian populations in our analysis had similar sample sizes and were painted using exactly the same procedure as the Jola and Fula, we do not see similar numbers of local ancestry changes in them, suggesting that such deviations are not linked to our chromosome painting procedure.

In addition to working with populations that have substantial recent admixture, there are further considerations to the results that come from our approach. The local ancestry inference procedure that we have used (Lawson et al., 2012) is based on a fully probabilistic framework. To access uncertainty in local ancestry with the paintings, we generated 10 realisations (sampled paintings) of the painting algorithm and inferred ancestry proportions from these paintings. Because of the stochastic nature of such an approach, we are unlikely to capture the full uncertainty in local ancestry, but chose this approach based on computational requirements and the use of a similar approach in previous admixture inference work (Hellenthal et al., 2014). Nevertheless, future work to optimise the computation issues of storing the *n * n* posterior copying probability matrix at each locus in the genome will likely provide a clearer assessment of local ancestry uncertainty. We used a binomial likelihood model, which takes each ancestry separately and models genome-wide proportions against local deviations. Using this model, we found no evidence that more than one ancestry increases or decreases in concert at a locus, at least in the Fula [Fig. 3]. Modelling changes across all ancestries at the same time, for example by using a multi-variate normal model, might provide further insights into part of the genome where we observe co-deviations.

## Conclusion

Gene-flow between closely related groups is increasingly being implicated as a key driver of adaptation. Here we have sought to address whether this process has resulted from admixture events which we previously inferred in African populations within the last 4,000 years. Our approach uncovered regions of the genome previously known to have undergone selection such as the *LCT/MCM6* gene complex in relation to lactose persistence, and *DARC* in relation to *P. vivax* malaria in the Fula of West Africa. In both cases, combining these observations with our understanding of the admixture history of the Fula demonstrates that changes in local ancestry can help pinpoint regions of the genome where natural selection may have been working. In addition, we identified other regions of the genome where local ancestry is deviates from expectations [Fig. 5], raising the possibility that adaptive gene-flow may be a more general, albeit rare, phenomenon. However, to evaluate how widespread this phenomenon is further work is needed to develop approaches that are able to detect more subtle signatures of ancestry deviations, and to account for the problems of non-independence highlighted above. Nevertheless, whilst adaptive introgression (i.e. adaptation resulting from interspecific gene-flow) has a deep literature, both theoretically (Lewontin and Birch, 1966) and empirically (for example in animals reviewed in Hedrick (2013)), examples of such an effect within species on a recent timescale are less common. As large genome-scale datasets from humans and other species become further available, the framework and methodology described here can be used to understand how widespread the phenomenon of adaptive gene-flow is.

## Acknowledgements

The data used for these analyses was released by MalariaGEN alongside the previous publication (see Busby et al., 2016). We thank all the MalariaGEN study sites that contributed samples to this analysis: a list of researchers involved at each study site can be found at https://www.malariagen.net/projects/host/consortiummembers. MalariaGEN is funded by the Wellcome Trust (WT077383/Z/05/Z, 090770/Z/09/Z) and the Bill and Melinda Gates Foundation through the Foundation for the National Institutes of Health (566).

Genotyping was performed at the Wellcome Trust Sanger Institute, partly funded by its core award from the Wellcome Trust (098051/Z/05/Z). This research was also supported by Centre grants from the Wellcome Trust (090532/Z/09/Z) and the Medical Research Council (G0600718). C.C.A.S. was supported by a Wellcome Trust Career Development Fellowship (097364/Z/11/Z).

## Materials and Methods

### Dataset

#### African population genetics dataset

We used the dataset of 3,283 individuals from 48 African and 12 Eurasian populations typed at 328,000 SNPs previously published by us (Busby et al., 2016), and available at https://www.malariagen.net/ resource/18. This dataset comprises 14 Eurasian and 46 African groups, mostly from sub-Saharan Africa [Fig. S1]. Half (23) of the African groups represent subsets of samples collected as part to the MalariaGEN consortium. Details of partners involved in the consortium and recruitment of samples in relation to studying malaria genetics are published elsewhere (Malaria Genomic Epidemiology Network, 2015). The remaining groups are from publicly available datasets and the 1000 Genomes Project, with Eurasian groups from the latter included to help understand the genetic contribution from outside of the continent. To study patterns of diversity, we created an integrated dataset at a common set of high-quality SNPs and used SHAPEITv2 (Delaneau et al., 2012) to haplotypically phase these genotype data, generating a final set of 6566 haplotypes.

### Defining Ancestry Regions

We used a combination of geographic, ethnic, and genetic information to group our sample into populations and eight ancestry regions [Fig. 1]. We analysed each population in turn, and inferred local ancestry using only those populations from outside of their ancestry region, which we term non-local paintings.

### Chromosome painting procedure

#### Inferring population specific prior copying probabilities and painting parameters

We utilised ChromoPainter (Lawson et al., 2012) to infer local ancestry. In order to do so we first needed to generate population-specific prior copying probabilities. The algorithm initially assumes that each donor haplotype is equally likely to be copied from, which in reality is not the case. Moreover, when there is less contextual information available for painting, for example at the beginning and ends of chromosomes, in its default setting, the algorithm will assign ancestry based on the on the initial prior copying probabilities, which will lead to noisy ancestry inference. We therefore performed initial runs of ChromoPainter where for each recipient population we used the Expectation-Maximisation (EM) algorithm to infer the prior copying probabilities from each recipient group (using the −*ip* flag). We ran ten EM iterations across 4 chromosomes (1,4,10,15) using 5 recipient individuals per population, and averaging across individuals and chromosomes to infer population specific prior copying probabilities.

We next estimated the recombination scaling constant, using the −*in* flag, and the mutation (HMM emission) probabilities, using the −*im* flag, separately for the same 5 individuals and 4 chromosomes noted above for each recipient population, and again averaged across chromosomes and individuals. We then painted all individuals from a given recipient population using these population-specific prior copying probabilities and model parameters.

#### Inferring local ancestry with ChromoPainter

To estimate local ancestry uncertainty, we take multiple sample paths through the HMM to generate 10 ancestry paintings per recipient haplotype. We note that in an ideal scenario, we would use ChromoPainter to infer the true uncertainty in the ancestry at a SNP by computing the full set of posterior marginal copying probabilities for each recipient haplotype at every position of the genome, giving us the probability that a recipient haplotype copies from all donor haplotypes at a given position. However, given *N* recipient haplotypes and *M* donor individuals, this would give an *N * M* matrix at each of *L* loci for each recipient population, which quickly leads to computational storage issues. To overcome this, we instead store 10 painting samples, leading to *N* * 10 matrix being stored at each locus and use these painting samples to estimate uncertainty in the local ancestry inference.

### Identifying local deviations in local ancestry

#### Binomial logistic model

The approach we take for identifying deviations in local ancestry is based on the idea that, in the absence of any other population genetic process, an individual’s ancestry proportions at a test SNP should be within the distribution of their genome-wide ancestry proportions. In theory, we can generate this null distribution of ancestry expectations based on the paintings at all genome-wide SNPs, or a subset thereof. In practice, we use a leave-one-out approach, and compute an individual’s genome-wide ancestry proportions from all SNPs that are not on the same chromosome as our test SNP that we want to test.

We model the proportion from each donor ancestry region separately via a binomial model with a mean estimated genome-wide and a linear increase on the logistic scale determined by β. We used a likelihood ratio test to determine if this parameter was significantly different from zero at its maximum.

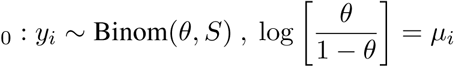

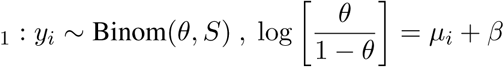

We calculate the MLE of beta by finding 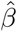 that solves:

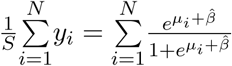

Then we use that MLE to calculate our likelihood ratio test statistic and compare it to a 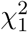 distribution to get our pvalue.

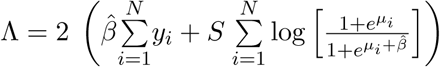

We then use chi-squared distribution to test if this statistic is in the tail.

## Supplementary Figures and Table

**Figure S1:**
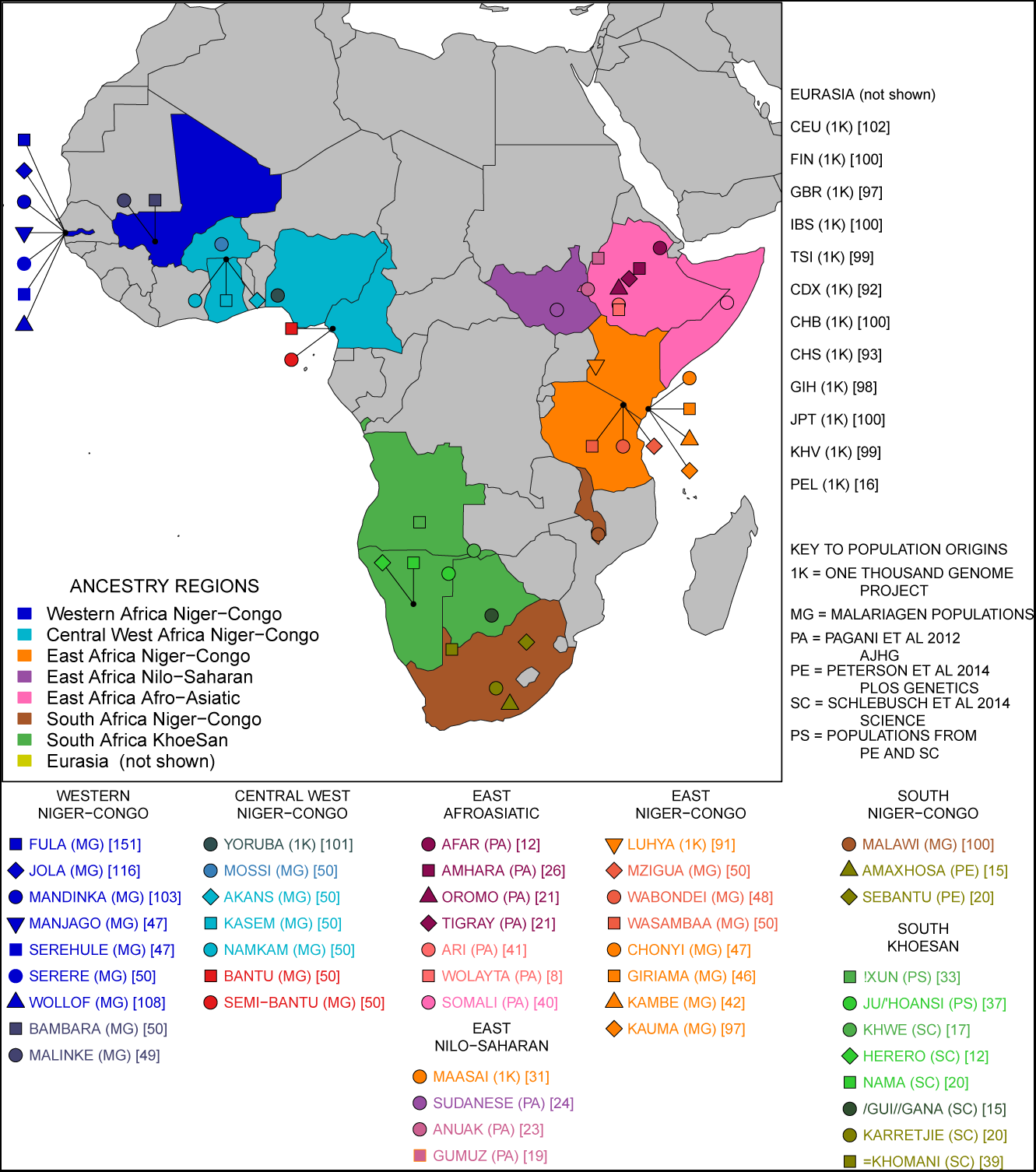
Description of population s used. Africa is divided into 7 ancestry regions based on ethnography, geography, and genetics.

**Figure S2:**
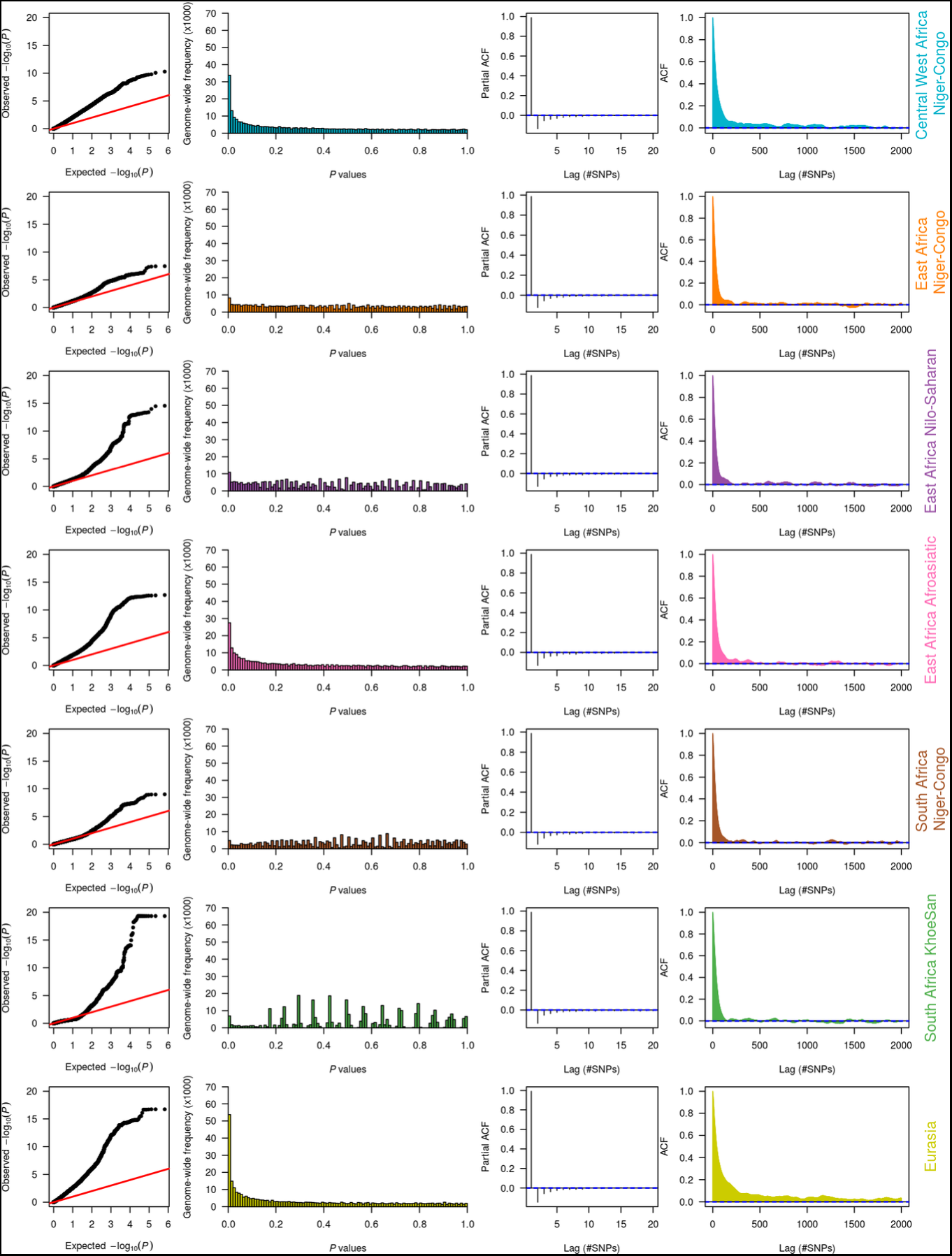
Quantile-Quantile plots of the model statistics in the Fula show inflation. We show QQ plots for *P* values generated by the binomial likelihood model for each ancestry separately.

**Figure S3:**
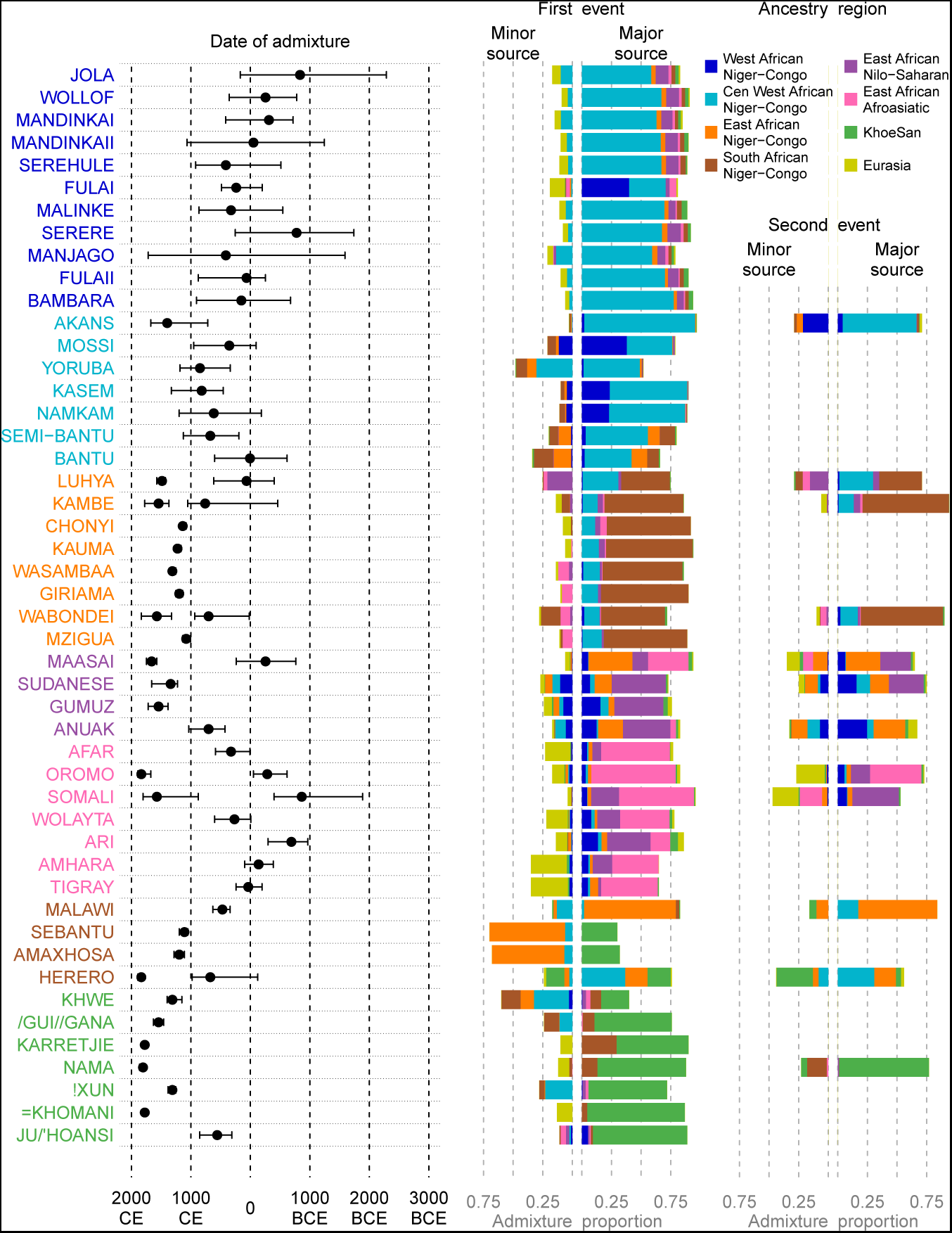
Overview of admixture events inferred in Sub-Saharan Africa. For reference, we show the main admixture events inferred previously by Busby et al. (2016). For each of 48 African populations, we characterise the main admixture event inferred from chromosome painting using *GLOBETROITER* (Hellenthal et al., 2014) including the date(s) of admixture and the ancestry composition of the two admixture sources contributing to an admixture event. In 12 populations we infer two admixture events which happened at the same time if there is a single admixture date, or, in the case of two events, reflect the source composition of the earlier event. In all cases the sum of the minor and major admixture sources bars adds up to 1

**Figure S4:**
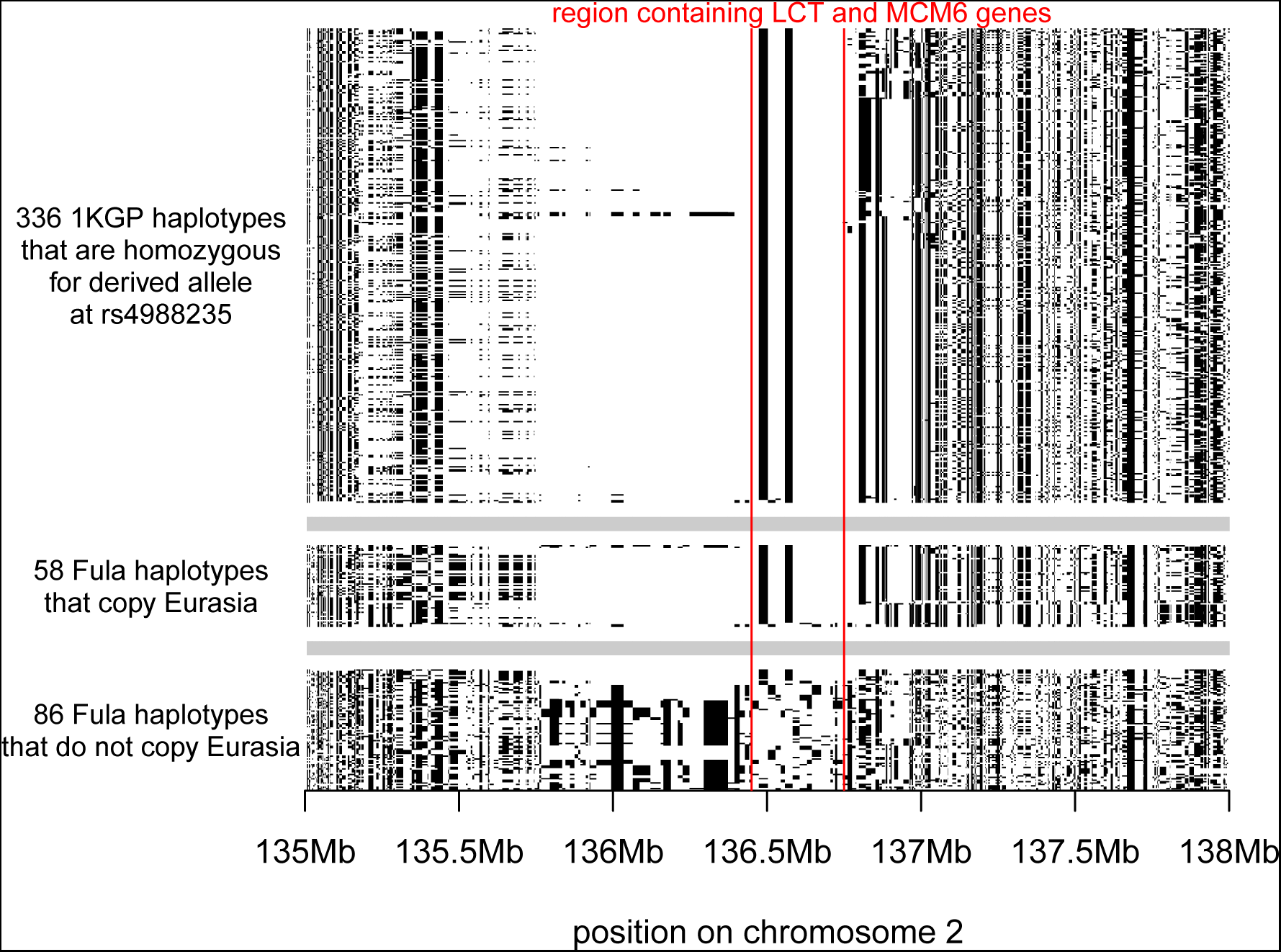
Comparison of Fula and One Thousand Genome Project (1KGP) haplotypes at the *LCT/MCM6* region. rs4988235 is one of the two main SNPs present in the lactase persistence haplotype. The SNP is not present in our dataset. To ascertain whether the signal we identified in this region with our method is indeed due to the presence of the derived allelle at this SNP, we show the phased haplotypes of all 1KGP Eurasian individuals who are homozygous for the derived variant at rs4988235 as well as the haplotypes for those Fula who copy Eurasia donors across the *LCT* region, and those that do not. The same haplotype is shared across the known 1KGP lactase persistence allele carriers and the Fula individuals who copy Eurasia, suggesting that the signal is indeed related to the presence of the lactase persistence haplotype in the Fula.

**Figure S5:**
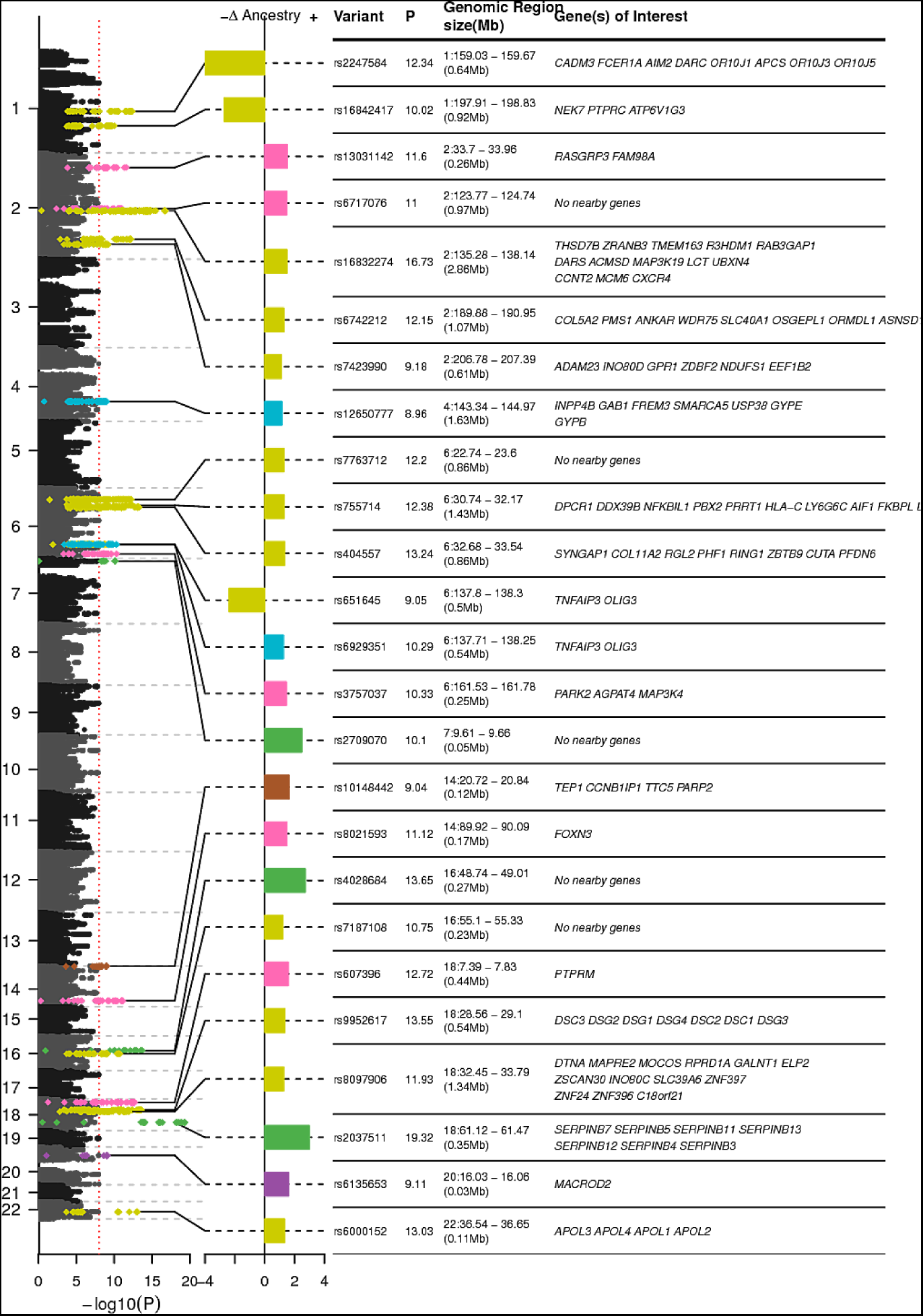
Manhattan plot showing ancestry deviations in the Fula. The left panel shows *−log*_10_ P values based on the binomial likelihood test for each of the 7 non-self ancestries tested in the Fula. SNPs with *−log*_10_P values greater than 8 are highlighted with the colour of the ancestry that the deviation is seen. The central panel shows the change (Δ) in ancestry observed at significant SNPs.The right panel gives information about the top ‘hit’ SNP and the genes within the genomic region around the hit.

**Figure S6:**
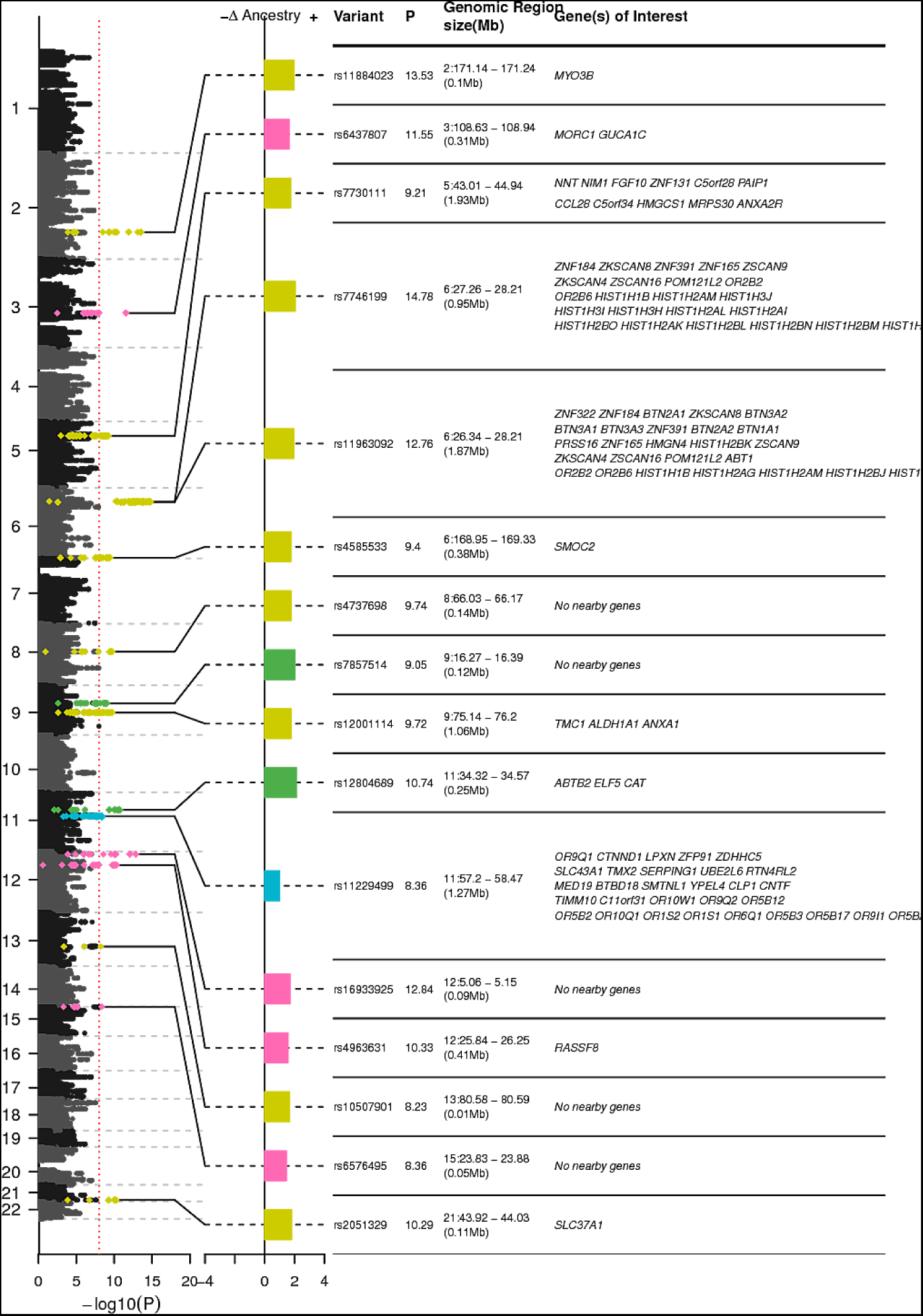
Manhattan plot showing ancestry deviations in the Jola. The left panel shows *−log*_10_ P values based on the binomial likelihood test for each of the 7 non-self ancestries tested in the Fula. SNPs with P values greater than 8 are highlighted with the colour of the ancestry that the deviation is seen. The central panel shows the change (Δ) in ancestry observed at significant SNPs.The right panel gives information about the top ‘hit’ SNP and the genes within the genomic region around the hit.

**Figure S7:**
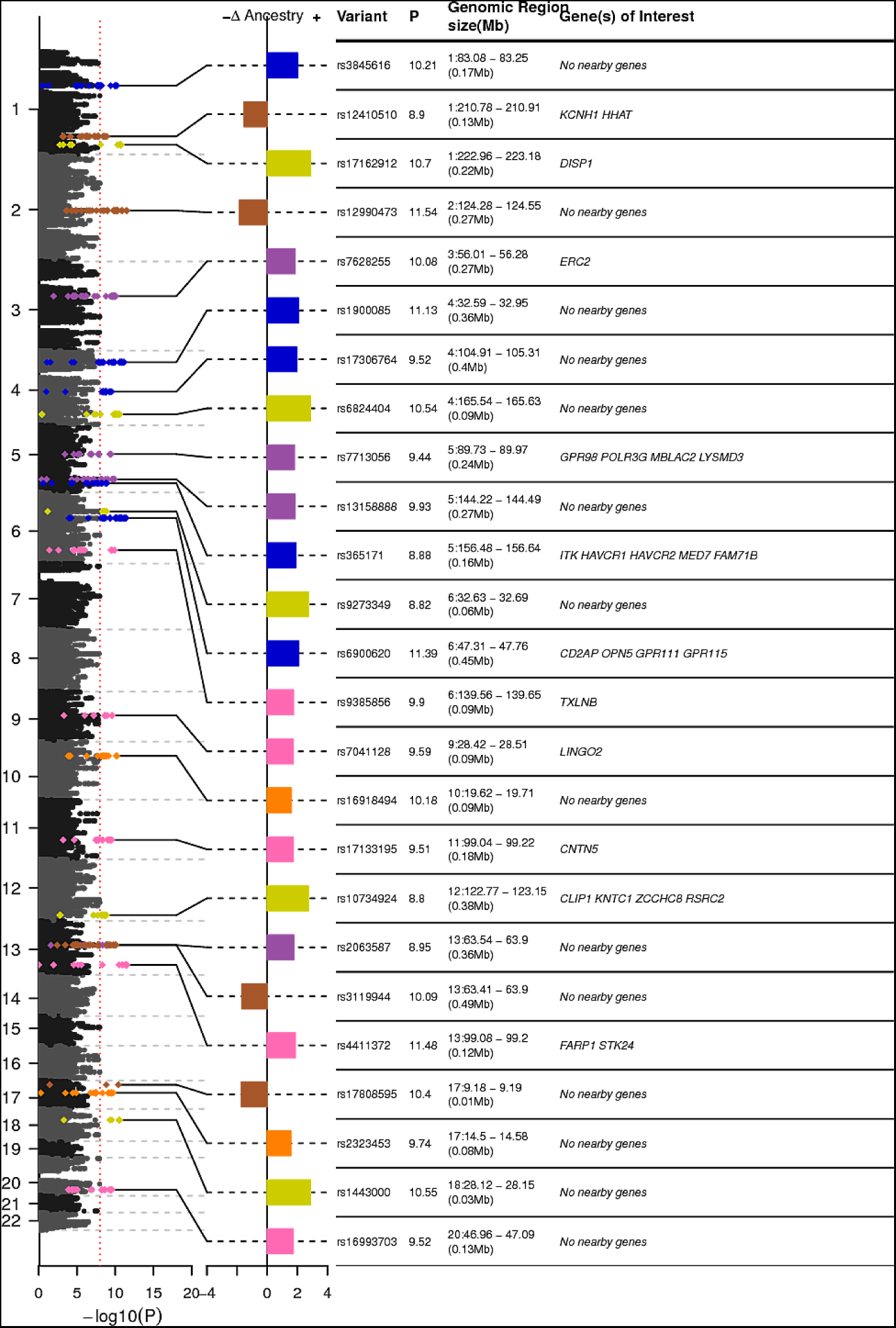
Manhattan plot showing ancestry deviations in the Ju/hoansi. The left panel shows *−log*_10_ P values based on the binomial likelihood test for each of the 7 non-self ancestries tested in the Fula. SNPs with P values greater than 8 are highlighted with the colour of the ancestry that the deviation is seen. The central panel shows the change (Δ) in ancestry observed at significant SNPs.The right panel gives information about the top ‘hit’ SNP and the genes within the genomic region around the hit.

**Figure S8:**
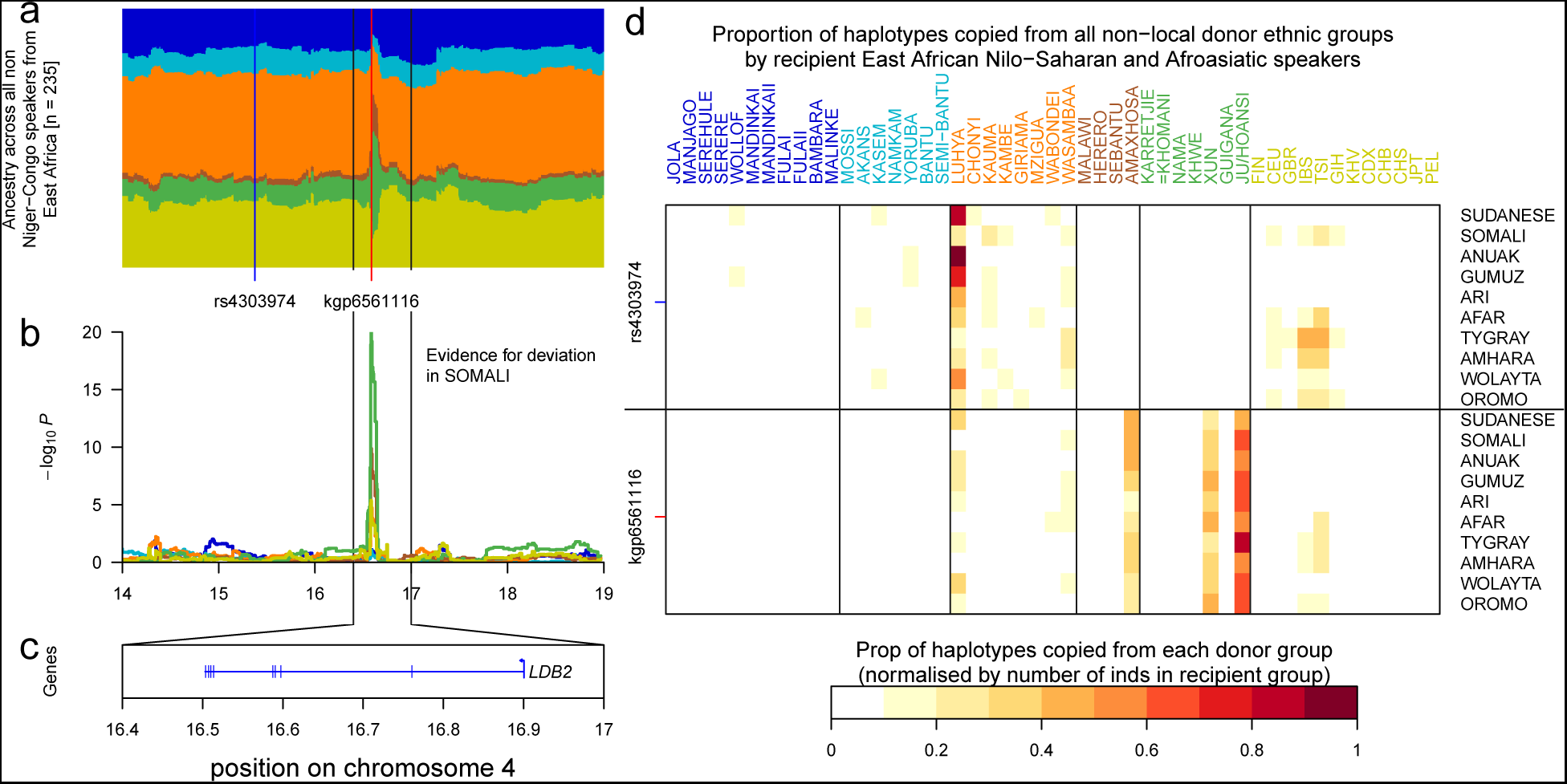
Increase d Khoesan ancestry at LDB2. **a** local ancestry across 4700 painted chromosomes from East African Nilo-Saharan and Afroasiatic speakers at the 14-19Mb region of chromosome 4. **b** results of the binomial ancestry deviation test shows a significant increase in Khoesan (green) ancestry between 16.2 and 16.8Mb. **c** UCSC genes across this region. **d** comparison of donor copying at two SNPs on chromosome 4, rs43003974 is 1.5Mb upstream of the main signal and most copying at this SNP comes from the Luhya; conversely at kgp656116 most copying comes from three Southern African groups: The Ju/hoansi, Xun, and Amaxhosa.

**Table S1:**
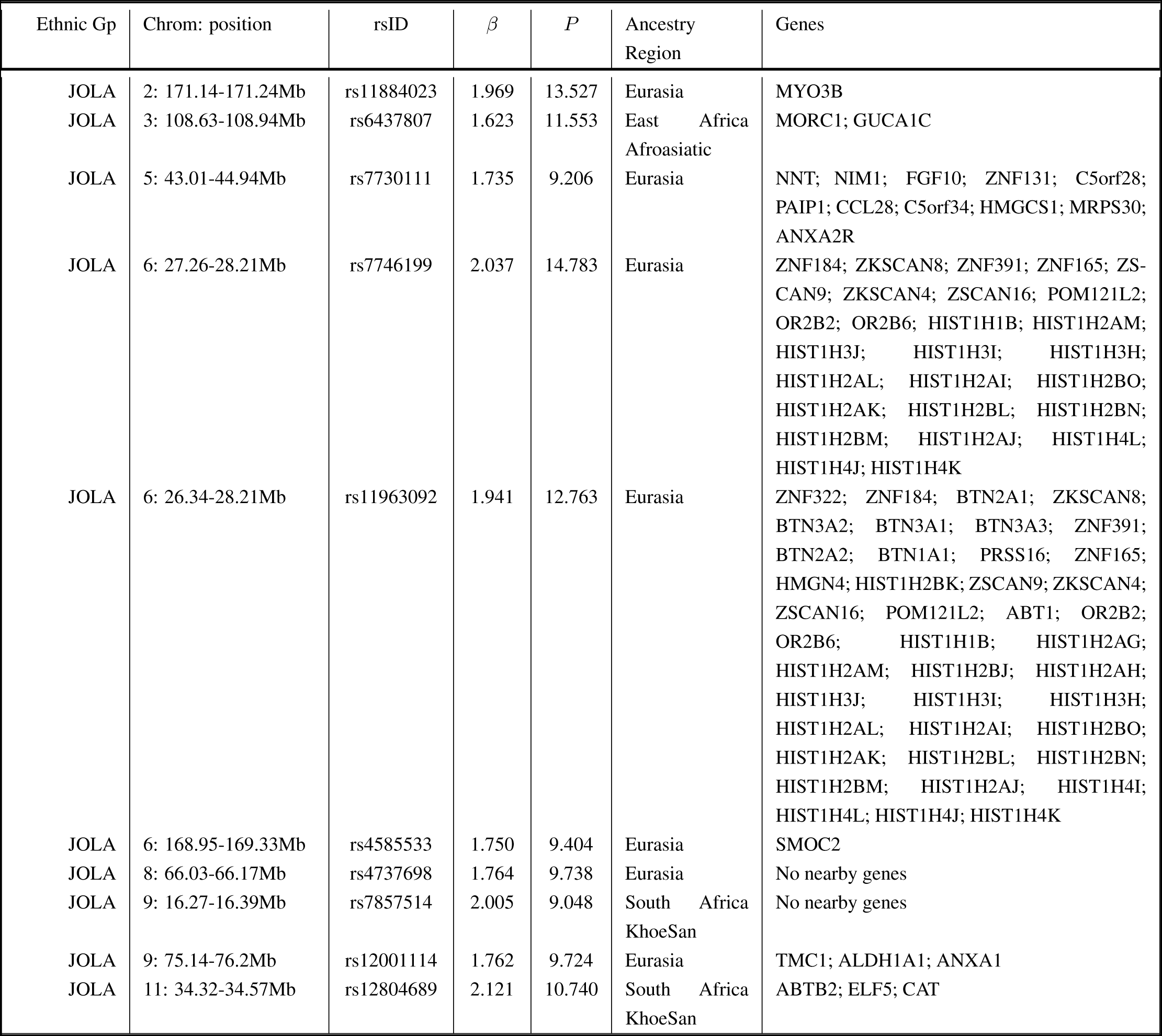
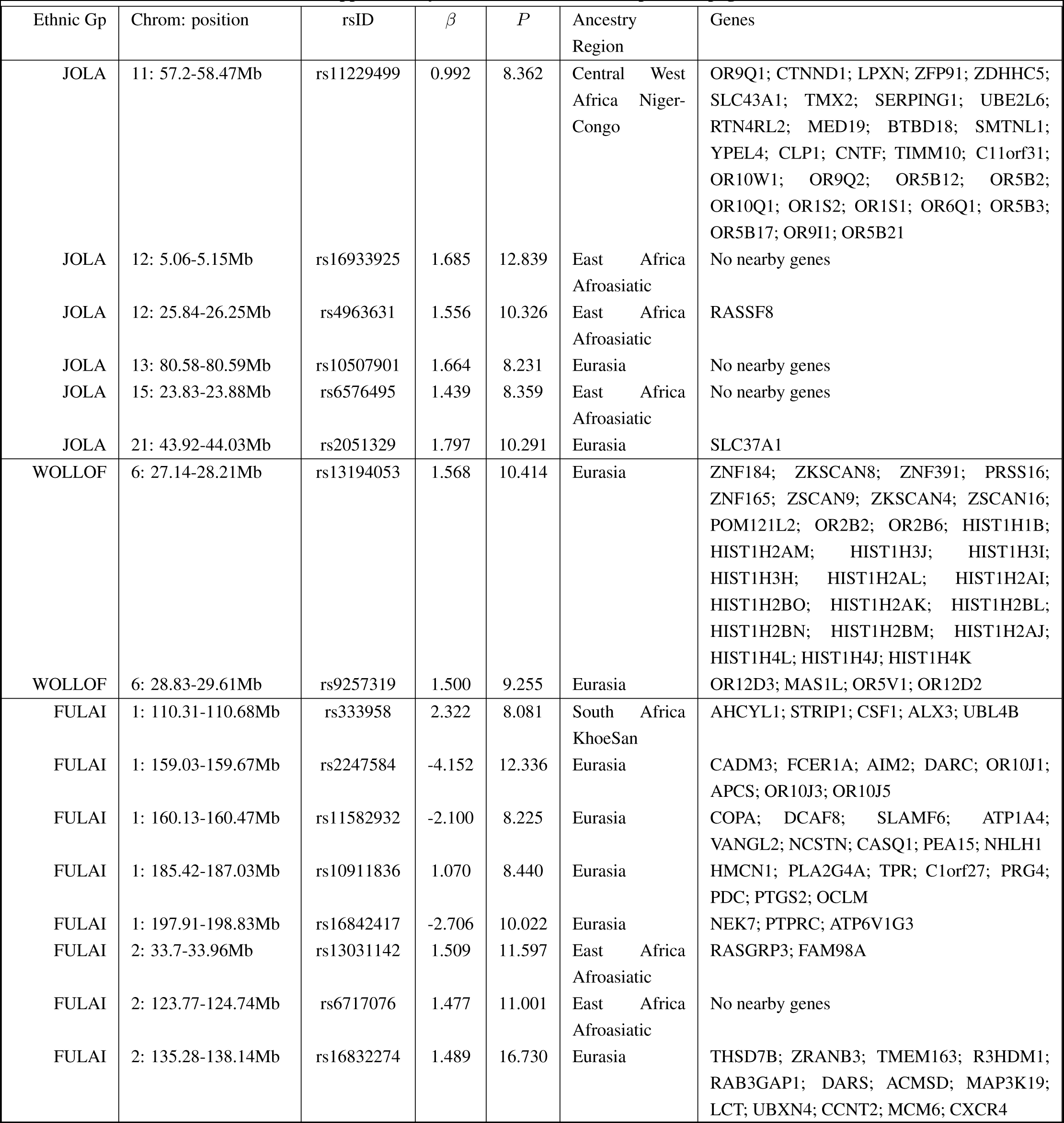
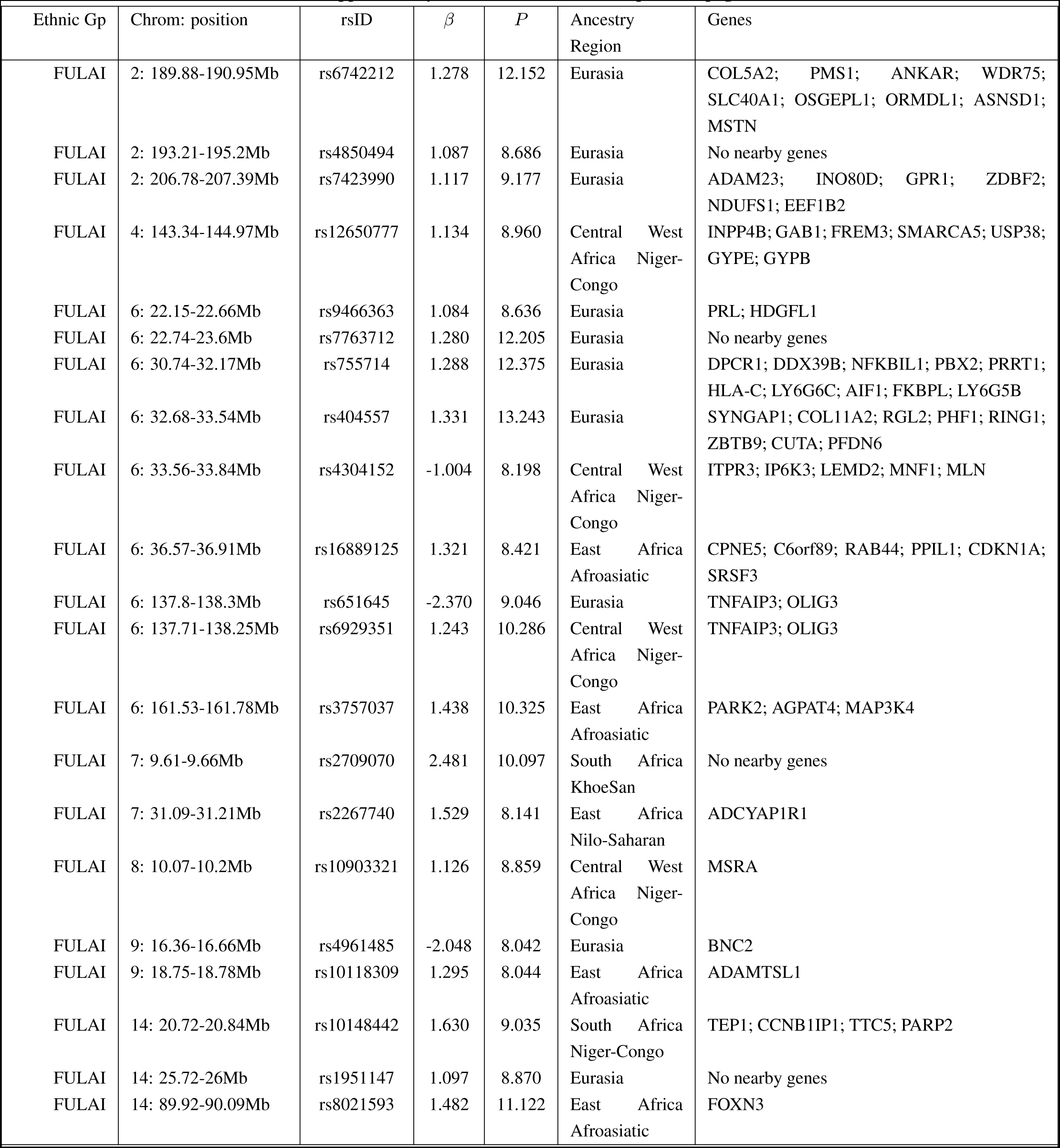
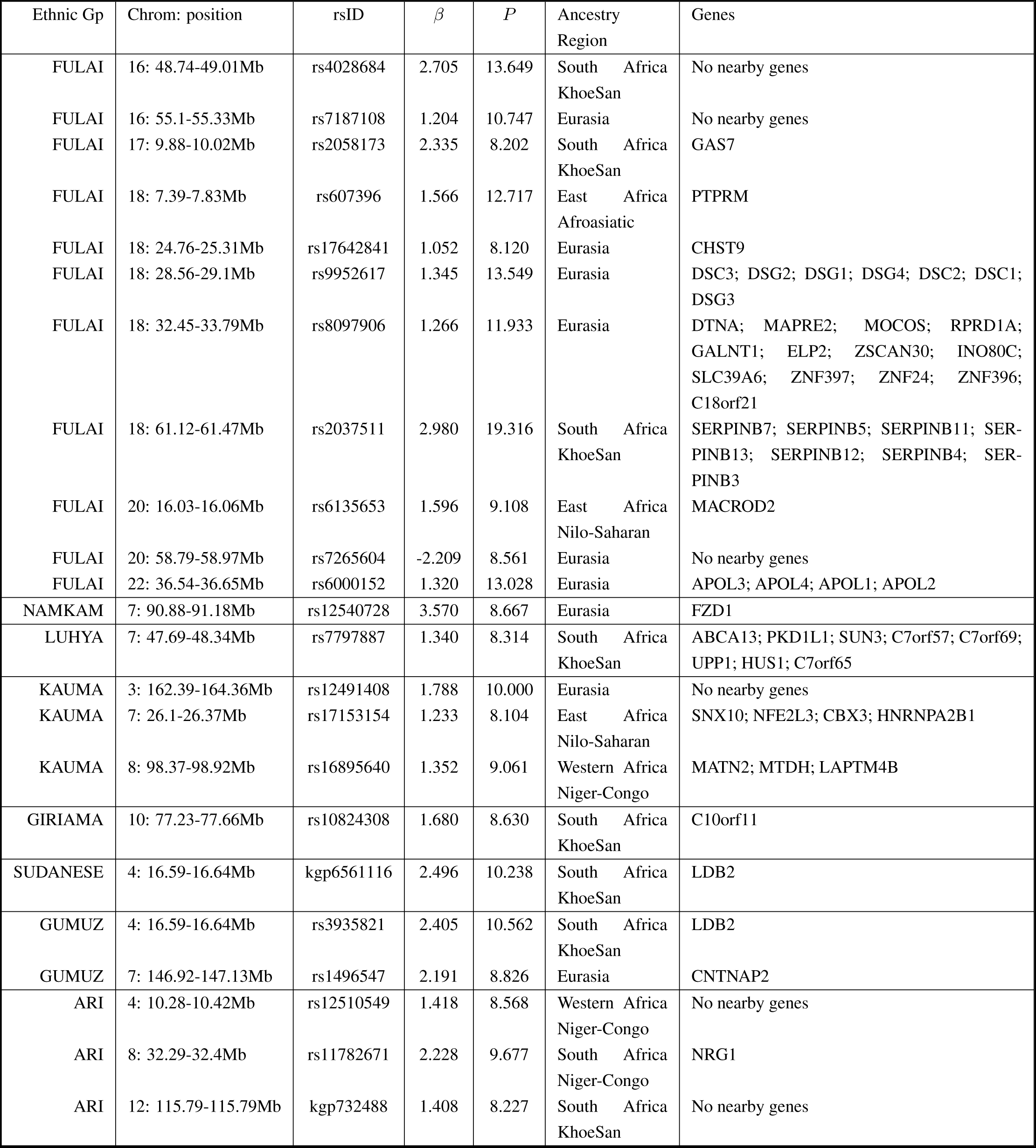
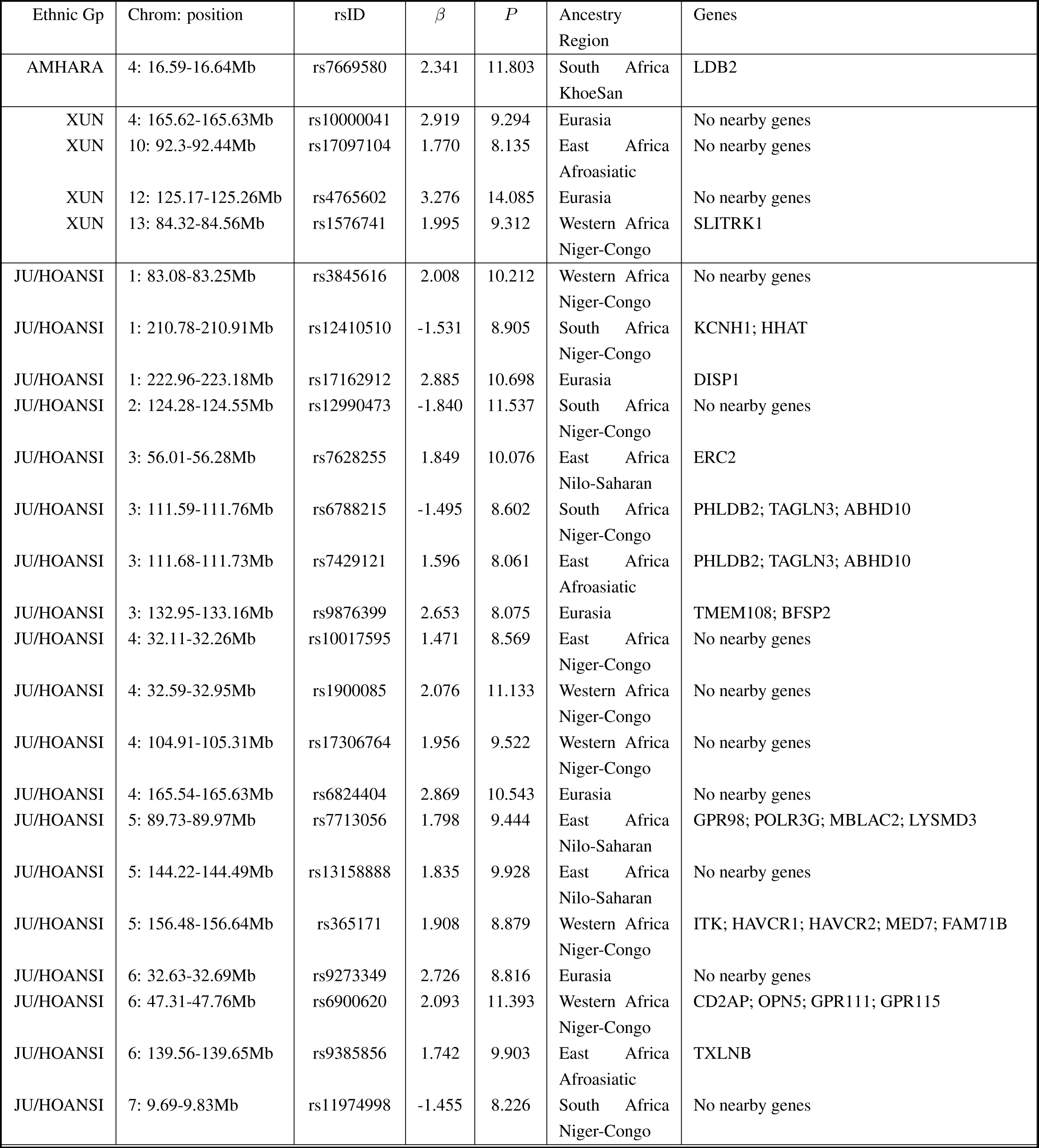
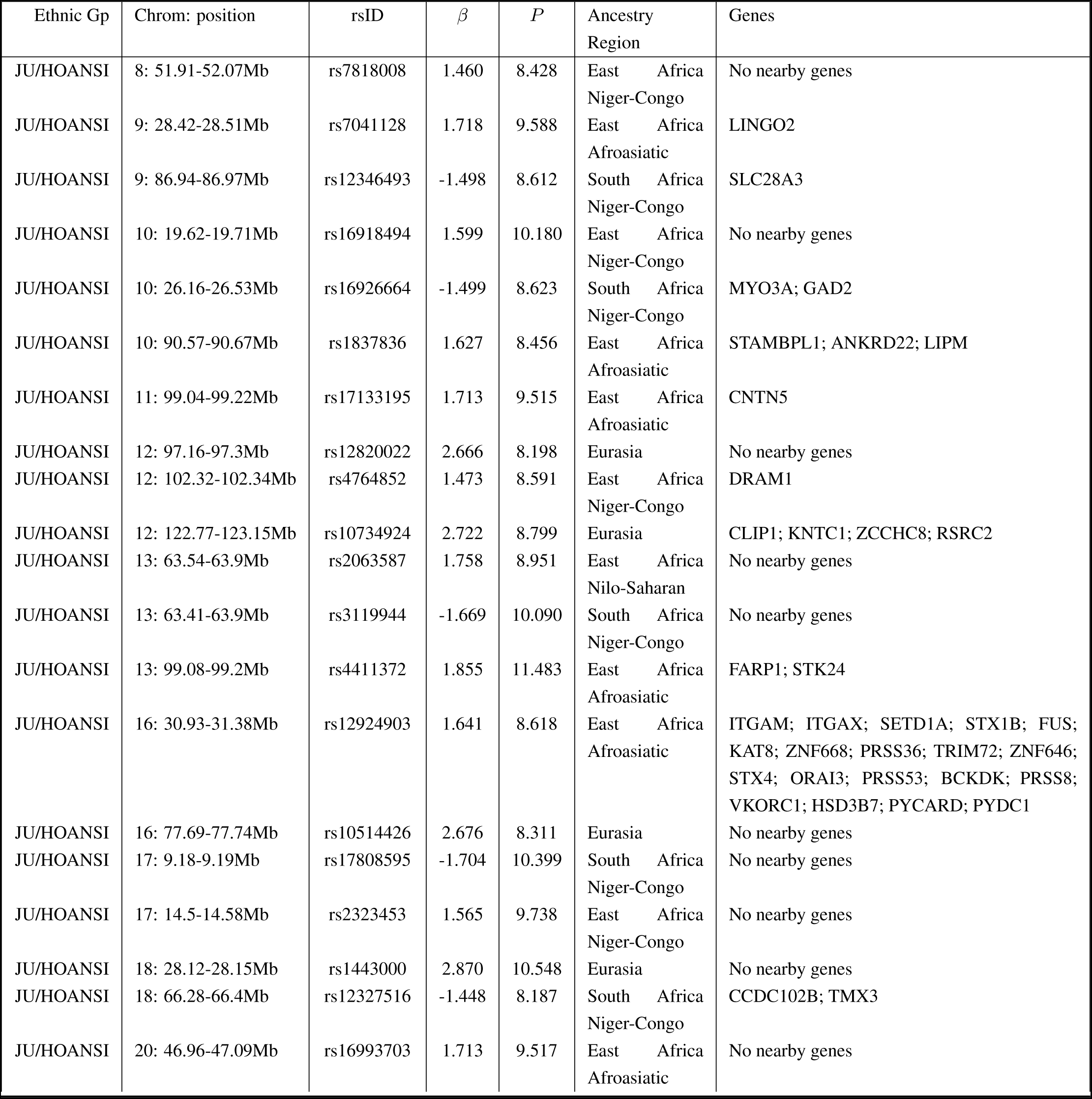
**Evidence for ancestry deviations in African populations.** Details of the ancestry deviation signals identified using the ancestry deviation model. We show the chromosome and position of the region where we observe a significant deviation in ancestry together with the rsID of the SNP with the largest deviation, the size and direction of the deviation, *β* and associated *P* value, the donor ancestry region of the deviation and the genes in the region of the signal for all significant signals of ancestry deviation.

